# Maternal-fetal interfaces transcriptome changes associated with placental insufficiency and a novel gene therapy intervention

**DOI:** 10.1101/2024.06.05.597595

**Authors:** Helen N. Jones, Baylea N. Davenport, Rebecca L. Wilson

**Affiliations:** Department of Physiology and Aging, College of Medicine, University of Florida, Gainesville, FL, USA; Center for Research in Perinatal Outcomes, College of Medicine, University of Florida, Gainesville, FL, USA

**Author notes:** Corresponding Author: Rebecca Wilson, Center for Research in Perinatal Outcomes, University of Florida, Gainesville, Florida 32610.

**Keywords:** trophoblast invasion, fetal growth restriction, placental insufficiency, nanoparticle, IGF1

## Abstract

The etiology of fetal growth restriction (FGR) is multifactorial, although many cases often involve placental insufficiency. Placental insufficiency is associated with inadequate trophoblast invasion resulting in high resistance to blood flow, decreased availability of nutrients, and increased hypoxia. We have developed a non-viral, polymer-based nanoparticle that facilitates delivery and transient gene expression of *human insulin-like 1 growth factor* (*hIGF1*) in placental trophoblast for the treatment of placenta insufficiency and FGR. Using the established guinea pig maternal nutrient restriction (MNR) model of placental insufficiency and FGR, the aim of the study was to identify novel pathways in the sub-placenta/decidua that provide insight into the underlying mechanism driving placental insufficiency, and may be corrected with *hIGF1* nanoparticle treatment. Pregnant guinea pigs underwent ultrasound-guided sham or *hIGF1* nanoparticle treatment at mid-pregnancy, and sub-placenta/decidua tissue was collected 5 days later. Transcriptome analysis was performed using RNA Sequencing on the Illumina platform. The MNR sub-placenta/decidua demonstrated fewer maternal spiral arteries lined by trophoblast, shallower trophoblast invasion and downregulation of genelists involved in the regulation of cell migration. *hIGF1* nanoparticle treatment resulted in marked changes to transporter activity in the MNR + *hIGF1* sub-placenta/decidua when compared to sham MNR. Under normal growth conditions however, *hIGF1* nanoparticle treatment decreased genelists enriched for kinase signaling pathways and increased genelists enriched for proteolysis indicative of homeostasis. Overall, this study identified changes to the sub-placenta/decidua transcriptome that likely result in inadequate trophoblast invasion and increases our understanding of pathways that *hIGF1* nanoparticle treatment acts on in order to restore or maintain appropriate placenta function.

## Introduction

FGR (estimated fetal weight <10^th^ percentile) occurs in up to 10% of pregnancies with suboptimal uteroplacental perfusion accounting for 25-30% of cases [1, 2]. Infants born FGR are at increased risk of perinatal morbidity and mortality, primarily due to prematurity and hypoxia [3]. Maintaining an upward growth trajectory *in utero* is paramount to preventing iatrogenic preterm delivery, neonatal intensive care unit (NICU) admission and associated complications. Additionally, the difficulties faced by the small fetus continue well beyond the perinatal period and across the entire lifespan [4-6].

The etiology of FGR is multifactorial, although many cases often involve placental insufficiency [7]. The placenta is a complex organ, essential for the transfer of nutrients and gases from mother to fetus, and the elimination of fetal waste products. Additionally, the placenta functions as an endocrine organ, actively synthesizing hormones, growth factors and cytokines [8]. In the human, maternal blood enters the intervillous space of the placenta via uterine arteries which undergo specialized transformation in the first half of pregnancy to become large diameter vessels with low resistance [9]. This transformative process is performed by the invasive trophoblast which migrate from anchoring villi, and is characterized by gradual loss of the arterial wall musculoelastic structure, and replacement with extravillous trophoblast [10]. Inadequate trophoblast invasion creates an environment with high resistance to blood flow, resulting in decreased availability of nutrients to the placenta, and increased hypoxia [11]. Despite the importance of vascular remodeling and trophoblast invasion to placenta development, the mechanistic processes are poorly understood.

In humans, trophoblast invasion occurs in the first half of pregnancy. Therefore, animal models are the only direct route to studying trophoblast invasion in vivo. Comparable animal models, however, are difficult to find. Non-human primates are the closest analogue, however not all species have comparable invasion, or are difficult to obtain and expensive to study. Guinea pigs offer a promising, cost-effective alternative as the guinea pig maternal-fetal interface demonstrates an invasive phenotype like that of humans, and a sub-placenta [12]. The sub-placenta is a distinct region of the maternal-fetal interface, not involved in exchange of nutrients/waste, which contains progenitors to the invasive trophoblast, analogous to the cell columns of the human placenta [13]. Additionally, maternal stress, and placental insufficiency can be induced non-invasively using a moderate maternal nutrient restricted (MNR) diet [14, 15]. The guinea pig MNR model of FGR is well characterized, resulting in reduced fetal weight from mid-pregnancy without infertility, or increased rates of fetal loss.

Despite accumulating knowledge linking inadequate trophoblast invasion with placental insufficiency and FGR, the prediction rate ranges from 12-47%, with more than half of FGR infants diagnosed after birth [16]. Therefore, an effective treatment for FGR needs to be capable of correcting fetal growth trajectories after the establishment of placental insufficiency and diagnosis of reduced in utero growth. Non-viral, polymeric nanoparticles have passed safety regulations and are successfully being used to deliver gene therapies in human cancer clinical studies [1]. Our prior studies demonstrate the use of a non-viral, polymer-based nanoparticle that facilitates transient (does not integrate into the genome) gene delivery specifically to trophoblast for the treatment of FGR [3-9]. We have successfully shown efficient nanoparticle uptake and increased *hIGF1* expression in human syncytiotrophoblast *ex vivo* [17], and *in vivo* using mice [3, 9], guinea pig [4, 5, 7, 8] and nonhuman primates [18]. More specifically in guinea pigs, we have shown efficient placental *hIGF1* nanoparticle uptake, and robust *hIGF1* expression leading to structural and functional changes in the placenta conducive of supporting fetal growth. Furthermore, we do not observe off-target expression of *hIGF1* nanoparticle in fetal tissues [19, 20].

Here, we assessed trophoblast invasion and the sub-placenta/decidua transcriptome at mid-pregnancy in our well-established MNR guinea pig model that leads to placental insufficiency and FGR. As the development of a treatment that targets placental insufficiency and corrects FGR is the ultimate goal, the aim of the study was to identify novel pathways in the sub-placenta/decidua that will provide insight into the underlying mechanism driving FGR and may be corrected with intervention.

## Materials and Methods

### Maternal Nutrient Restriction Model of Fetal Growth Restriction

Animal care and usage was approved by the Institutional Animal Care and Use Committees at Cincinnati Children’s Hospital and Medical Center (Protocol number 2017-0065) and the University of Florida (Protocol number 202011236). Details of maternal nutrient restriction model, and ultrasound-guided intra-placental *hIGF1* nanoparticle injections have been published [14, 15, 21]. Female (dams) Dunkin-Hartley guinea pigs were purchased (Charles River Laboratories, Wilmington, MA) at 500-550 g and housed individually in a temperature-controlled facility with a 12 h light-dark cycle. At initiation of maternal nutrient restriction (MNR), dams were weighed and systematically assigned to either the Control diet group (n = 6) or MNR diet group (n = 7). Control dams were provided food (Lab Diet 5025) and water was ad libitum; MNR dams were provided water ad libitum however, food intake was restricted to 70% per kilogram body weight of the Control group from at least four weeks preconception through to mid-pregnancy (GD30), thereafter increasing to 90% [14, 15]. At GD30-33, dams underwent an ultrasound-guided, transuterine, intra-placental injection of either sham (200 µL of PBS: Control n = 3 and MNR n = 3) or *hIGF1* nanoparticle (60 µg plasmid in 200 µL injection: Control + *hIGF1* n = 3 and MNR + *hIGF1* n = 4), as provided in detail [21]. Dams were sacrificed five (GD35-38) days after *hIGF1* nanoparticle treatment. Sub-placenta/decidua tissue from two fetuses per litter (one female and one male) were blunt dissected away from the rest of the maternal-fetal interface and snap-frozen in liquid nitrogen for later RNA extraction.

### Total RNA Isolation and RNA-Seq Library Preparation

Total RNA was extracted from frozen sub-placenta/decidua (Control n = 6; Control + *hIGF1* n = 3; MNR n = 6; MNR + *hIGF1* n = 8) using Qiagen RNeasy Midi extraction kits following the manufacturer’s instructions. RNA integrity numbers greater than 5 were used for RNA-Seq studies. RNA-Seq Libraries for each gender and experimental group were generated from 1.5 µg RNA using the Illumina Stranded mRNA Prep Kit by University of Florida Interdisciplinary Center for Biotechnology Research (ICBR) Gene Expression and Genotyping Core.

### RNA-Seq and Gene Expression Analysis

RNA-Seq Libraries were sequenced using the Illumina NovaSeq platform. Short reads were trimmed using trimmomatic (v 0.36) [22], and Quality Control on the original and trimmed reads was performed using FastQC (v 0.11.4) [23] and MultiQC [24].

The reads were aligned to the guinea pig transcriptome using STAR (v 2.7.9a) [25], and transcript abundance was quantified using RSEM (v 1.3.1) [26]. Differential expression analysis was performed using DESeq2 [27], with a raw P-value threshold ≤0.05 and Log2-fold change of ≤-1.0 and ≥1.0. Results report protein-coding genes as well as other transcript types. Raw and processed sequencing data is available on NCBI Geo (GSE269098).

Differentially expressed genes between different group comparisons were separated into upregulated genes (positive log changes) and downregulated genes (negative log changes) and entered into ToppFun (ToppGene Suite V31 [28]) for enrichment analysis of GO Biological Processes and GO Molecular Functions. P values were calculated using the Hypergeometric Probability Mass Function and false discovery rate corrected using Benjamini–Hochberg methods.

### Immunohistochemistry

Immunohistochemistry (IHC) was used to assess extent of trophoblast invasion [12, 13, 29] in the guinea pig sub-placenta/decidua (n = 4 placentas per group). 5 µm thick placenta sections were de-waxed and rehydrated following standard protocols. No antigen retrieval was performed. Slides were incubated in 3% hydrogen peroxide for 10 mins to block endogenous peroxidase activity. Serum-free protein block (*Dako*) was then applied for 30 min at room temperature, followed by incubation in primary antibodies, diluted in 10% goat serum, 1% BSA in PBS, for 30 min at 37ºC. Primary antibodies against cytokeratin (*Bethyl* A500-019A; 1:50) were used to visualize maternal vessels which had been colonized by trophoblast, and against SMA for smooth muscle actin (SMA: *Dako* M085129; 1:100) to visualize maternal vessels devoid of trophoblast. Following incubation with primary antibodies, sections were washed and then biotinylated anti-mouse IgG (*Vector* BA-9200; 1:500) secondary antibodies applied for 1 h at room temperature. Staining was amplified using the Vector ABC kit (Vector), and detected using DAB (Vector) for brown precipitate. Hematoxylin was used to counterstain nuclei. All sections were imaged using the Axioscan scanning microscope (*Zeiss*). The Zen Imaging software (*Zeiss*) was used to capture representative images and for invasion analysis. Distance of vessels (minimum 20 vessels per section), with cells either positive for cytokeratin or positive for SMA were calculated by drawing a line from the middle of the vessel lumen to the decidual edge of the section. Thickness of the sub-placenta/decidua region was also calculated by averaging 10 measurements from the edge of the sub-placenta cell columns to the decidual edge.

## Results

### Maternal nutrient restriction is associated with decreased number of trophoblast invaded maternal vessels and shallower invasion

To assess the impact of MNR on trophoblast invasion in the sub-placenta/decidua, vessels lined with trophoblast cells in the sub-placenta/decidua were visualized with immunohistochemistry and compared to vessels lined with endothelial cells expressing smooth muscle actin (Figure 1A). At mid-pregnancy, the number of vessels invaded by trophoblast (Figure 1A) and the depth of invasion (Figure 1B) was reduced with MNR compared to Control. *hIGF1* nanoparticle treatment did not affect either parameter and sub-placenta/decidua depth was similar across all groups (estimated marginal mean and 95% CI (µm): Control = 3190, 3045-3343; Control + *hIGF1* = 3230, 2909-3586; MNR = 3170, 2882-3488; MNR + *hIGF1* = 3028, 2626-3493, P>0.05).

**Fig. 1.**
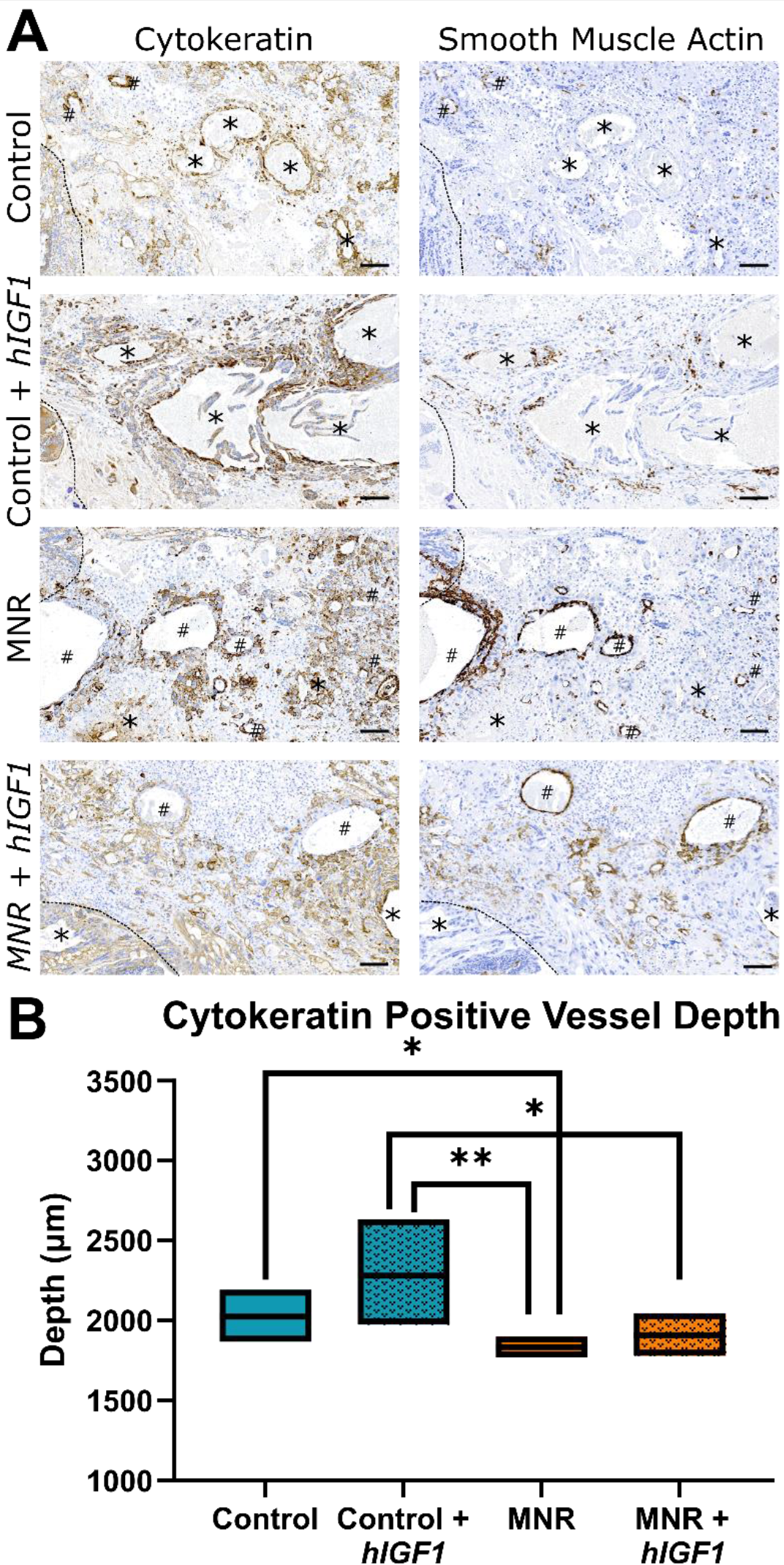
Representative serial sections of guinea pig sub-placenta/decidua stained for cytokeratin (trophoblast cells) and smooth muscle actin (SMA: endothelial cells), and depth of trophoblast invasion. **A**. In Control and Control placentas treated with *hIGF1* nanoparticle (Control + *hIGF1*) there were more vessels lined with trophoblast (Asterix) compared to uninvaded vessels represented by smooth muscle actin (SMA) lined vessels (hashtag). In maternal nutrient restricted (MNR) and MNR placentas treated with *hIGF1* nanoparticle (MNR + *hIGF1*), whilst there was clear trophoblast invasion, the number of vessels lined with trophoblast and not lined with SMA positive endothelial cells was diminished. **B**. Irrespective of *hIGF1* nanoparticle treatment, depth of vessels invaded with trophoblast was reduced with MNR compared to Control. n = 4-5 placentas stained per group. Data are estimated marginal mean ± 95% confidence interval. Dashed line marks edge of the sub-placenta trophoblast cell columns

### Maternal nutrient restriction and hIGF1 nanoparticle treatment alters transcriptome profiles in sub-placenta/decidua

Principal component analysis (PCA) on normalized expression data between sub-placenta/decidua tissue from female and male fetuses showed no strong confounding impact of fetal sex (Figure 2). Therefore, data generated from female and male sub-placenta/decidua were combined for further analysis. Initially, using an FDR (q-value) ≤ 0.05, very few transcripts were differentially expressed in sub-placenta/decidua from MNR dams compared to Control. Therefore, transcripts with a Log2 fold-change of ≤-1.0 or ≥1.0, and raw p-value of ≤0.05 were assessed for enrichment analysis (Figure 3). The full list of differentially expressed genes can be found in Supplemental Material.

**Fig. 2.**
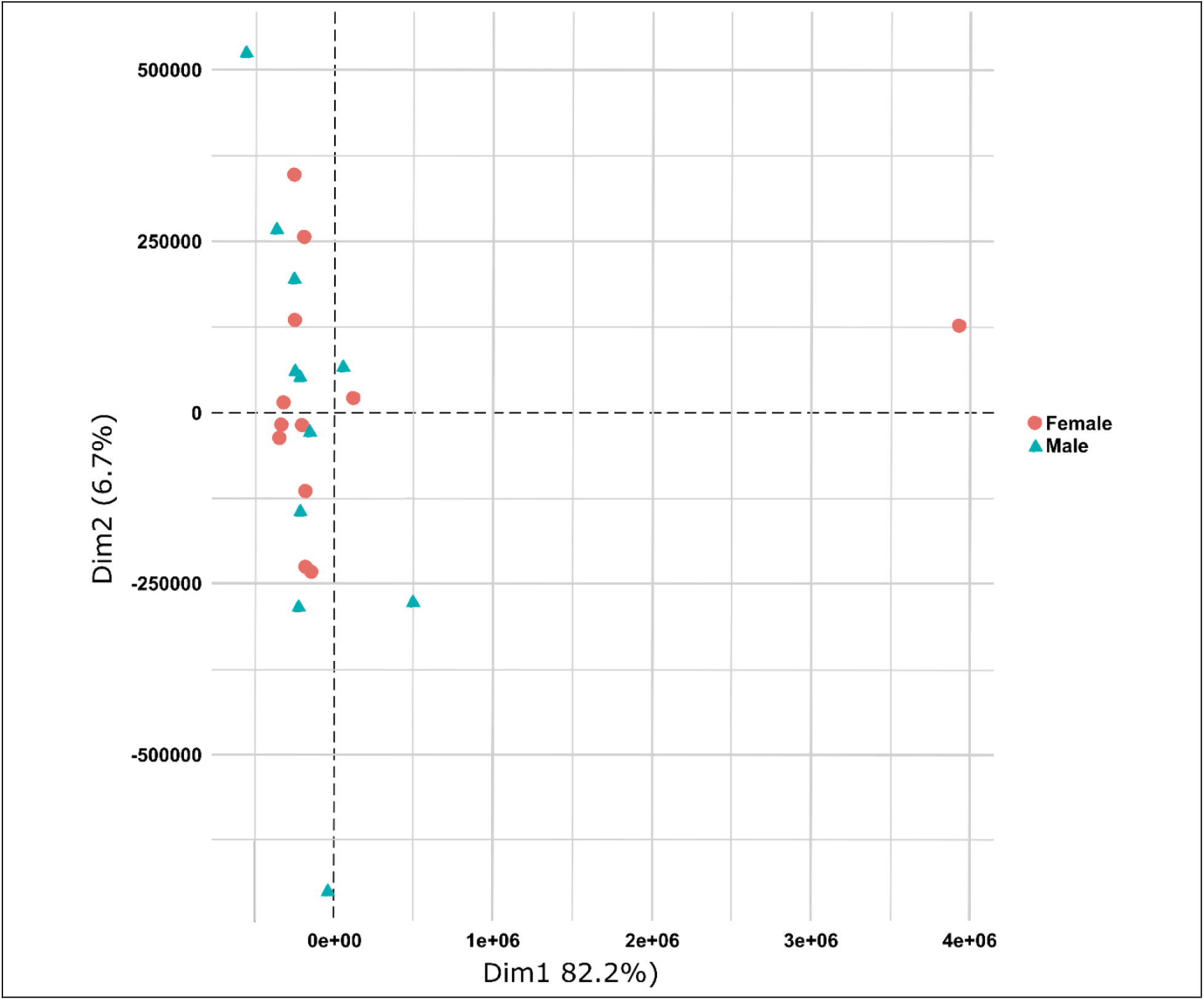
Principal component analysis (PCA) on normalized expression data between sub-placenta/decidua tissue from female and male fetuses. Normalized gene expression was not different enough between sub-placenta/decidua tissue of female and male fetuses to separate into distinct groups. n = 12 female and 11 male.

**Fig. 3.**
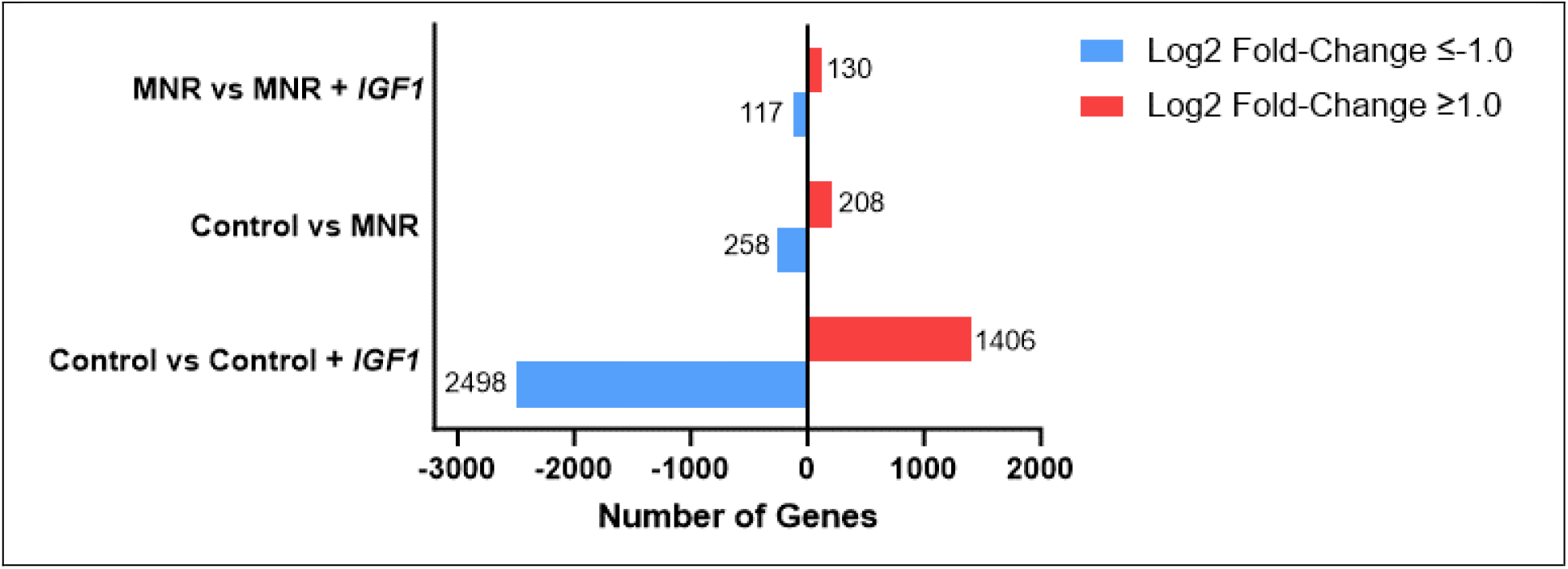
Number of differentially expressed genes in the guinea pig sub-placenta/decidua, modified by maternal nutrient restriction (MNR) and/or *hIGF1* nanoparticle treatment. Compared to Control, *hIGF1* nanoparticle treatment (Control + *hIGF1*) resulted in 2498 genes being downregulated and 1406 genes upregulated. MNR resulted in 258 genes being downregulated and 208 genes upregulated when compared to Control. In MNR sub-placenta/decidua, *hIGF1* nanoparticle treatment (MNR + *hIGF1*) resulted in 117 downregulated genes and 130 upregulated genes when compared to sham treated MNR. n = 5 Control, 3 Control + *hIGF1*, 6 MNR and 8 MNR + *hIGF1* sub-placenta/decidua samples.

### Maternal nutrient restriction is associated with reduced epithelium development, but hIGF1 nanoparticle treatment increases transporter activity

Previously, we have reported that MNR results in reduced ERK signaling activity, and increased expression of tight-junction/adhesion proteins in the sub-placenta/decidua [30]. Consistent with these previous findings, pathway enrichment analysis of genelists decreased in the MNR sub-placenta/decidua compared to Control revealed processes including Epithelium Development (FDR: 2.27E-02), Tube Morphogenesis (FDR: 3.92E-02), and Regulation of Cell Migration (FDR: 4.77E-02) (Table 1 and Supplemental Material). Genes in these processes included *Map3K1, Fgf10*, and *Cxcl10*.

**Table 1.** Pathway enrichment analysis of differentially expressed genes that were decreased in sham treated maternal nutrient restricted (MNR) sub-placenta/decidua compared to Control.

| ID | Pathway | p-value | FDR | Hit Count in Query List | Hit Count in Genome |
| --- | --- | --- | --- | --- | --- |
| GO:0060429 | epithelium development | 4.57E-05 | 2.27E-02 | 23 | 1979 |
| GO:0035239 | tube morphogenesis | 1.84E-04 | 3.92E-02 | 18 | 1467 |
| GO:0030334 | regulation of cell migration | 2.72E-04 | 4.77E-02 | 16 | 1249 |
| GO:0030198 | extracellular matrix organization | 1.14E-04 | 2.98E-02 | 9 | 394 |
| GO:0007584 | response to nutrient | 2.45E-04 | 4.77E-02 | 8 | 344 |
| GO:0034699 | response to luteinizing hormone | 2.26E-05 | 1.45E-02 | 3 | 12 |

Conversely, when the MNR placenta was treated with *hIGF1* nanoparticle (MNR + *hIGF1*), genelists increased in the sub-placenta/decidua were enriched for pathways including Transporter Activity (FDR: 4.89E-02), and Metallopeptidase Activity (FDR: 2.82E-02) when compared to sham-treated MNR sub-placenta/decidua (Table 2 and Supplemental Material). Increased nutrient transporter expression has been previously shown in MNR + *hIGF1* placentas [21]. In this study, increased genes in the MNR + *hIGF1* sub-placenta/decidua included *Nos1, Slc1A3*, and *Slc7A11*.

**Table 2.** Pathway enrichment analysis of differentially expressed genes increased in *hIGF1* nanoparticle treated maternal nutrient restricted (MNR) sub-placenta/decidua compared to sham treated MNR.

| ID | Pathway | p-value | FDR | Hit Count in Query List | Hit Count in Genome |
| --- | --- | --- | --- | --- | --- |
| GO:0005215 | transporter activity | 4.24E-03 | 4.89E-02 | 12 | 1427 |
| GO:0015075 | ion transmembrane transporter activity | 3.08E-03 | 4.39E-02 | 10 | 1017 |
| GO:0046873 | metal ion transmembrane transporter activity | 3.68E-03 | 4.39E-02 | 7 | 559 |
| GO:0046943 | carboxylic acid transmembrane transporter activity | 5.66E-04 | 2.25E-02 | 5 | 189 |
| GO:0005342 | organic acid transmembrane transporter activity | 5.80E-04 | 2.25E-02 | 5 | 190 |
| GO:0008237 | metallopeptidase activity | 1.10E-03 | 2.82E-02 | 5 | 219 |

### Placental hIGF1 nanoparticle treatment in control sub-placenta/decidua downregulates genelists related to transferase and kinase activity

In the Control + *hIGF1* sub-placenta/decidua, genelists that were decreased with *hIGF1* nanoparticle treatment were enriched for molecular functions including Adenyl Ribonucleotide Binding (FDR: 9.68E-07), ATP binding (FDR: 4.46E-07), Transferase Activity (FDR: 2.86E-05), and Kinase Activity (FDR: 3.29E-04) when compared to Control sham (Table 3). 152 genes comprising the Transferase Activity molecular function were reduced in Control + *hIGF1* sub-placenta/decidua, and included genes *Jak2, MapK13, Kdr, Nos2, Igf2, IgfR1*, and various *Pik3* isoforms. Other molecular functions enriched because of *hIGF1* nanoparticle treatment in the Control + *hIGF1* sub-placenta/decidua are listed in Supplementary Material. Genelists increased in the Control + *hIGF1* sub-placenta/decidua when compared to sham-treated Control were enriched for biological processes, including Proteolysis (FDR: 2.04E-02), Carbohydrate Derivative Metabolic Process (FDR: 1.56E-04), and Negative Regulation of Molecular Function (FDR: 1.28E-02)(Table 3). 80 genes involved in Negative Regulation of Molecular Function were increased, including *Nfkbia, Sfn, Pkn1, Rsp20*, and *Bcl2L1*.

**Table 3.** Pathway enrichment analysis of differentially expressed genes in *hIGF1* nanoparticle treated Control sub-placenta/decidua compared to sham treated Control.

| ID | Pathway | p-value | FDR | Hit Count in Query List | Hit Count in Genome |
| --- | --- | --- | --- | --- | --- |
| Decreased |  |  |  |  |  |
| GO:0032559 | adenyl ribonucleotide binding | 9.39E-10 | 9.68E-07 | 192 | 1606 |
| GO:0005524 | ATP binding | 2.17E-10 | 4.46E-07 | 187 | 1526 |
| GO:0016772 | transferase activity, transferring phosphorus-containing groups | 1.39E-08 | 7.14E-06 | 152 | 1242 |
| GO:0016301 | kinase activity | 7.98E-07 | 3.29E-04 | 128 | 1072 |
| GO:0051020 | GTPase binding | 5.33E-04 | 4.23E-02 | 46 | 356 |
| GO:0017147 | Wnt-protein binding | 1.80E-05 | 4.11E-03 | 11 | 32 |
| Increased |  |  |  |  |  |
| GO:0065003 | protein-containing complex assembly | 8.99E-06 | 1.37E-03 | 106 | 1859 |
| GO:0006508 | proteolysis | 2.09E-04 | 2.04E-02 | 105 | 1987 |
| GO:1901135 | carbohydrate derivative metabolic process | 7.10E-07 | 1.56E-04 | 82 | 1256 |
| GO:0044092 | negative regulation of molecular function | 1.19E-04 | 1.28E-02 | 80 | 1401 |
| GO:0051336 | regulation of hydrolase activity | 4.78E-04 | 3.73E-02 | 67 | 1178 |
| GO:0030162 | regulation of proteolysis | 4.72E-05 | 6.51E-03 | 57 | 880 |
| GO:0046034 | ATP metabolic process | 1.02E-08 | 1.20E-05 | 31 | 256 |

## Discussion

Using the guinea pig model, in which the sub-placenta is an analogous structure to the cell columns in humans and source of invasive trophoblast, we show that increased maternal stress and placental insufficiency is associated with fewer maternal spiral arteries lined by trophoblast and reduced trophoblast invasion in the MNR sub-placenta/decidua at mid-pregnancy which was associated with downregulation of genes involved in epithelial development and the regulation of cell migration. Additionally, short-term *hIGF1* nanoparticle treatment results in marked changes to transporter activity in the MNR + *hIGF1* sub-placenta/decidua indicative of increased capacity to transport nutrients. Under normal growth conditions however, *hIGF1* nanoparticle treatment in Control + *hIGF1* is associated with downregulation of kinase signaling and increased proteolysis indicative of homeostasis.

In early human pregnancy, conversion of maternal spiral arteries into larger, more competent vessels is crucial for normal placentation [9]. Failure to adequately remodel the maternal spiral arteries is a key characteristic of pregnancy complications and thus is a major focus of placental research. In vivo investigations of trophoblast invasion in the humans is not available due to ethical reasons, hence animal models are used as a surrogate. Extensive investigations have been performed showing the similarities in trophoblast invasion between humans and caviomorphs including guinea pigs [13, 31]. As with humans, invasive trophoblasts in the guinea pig originate from clusters of non-invasive, proliferating stem cells located in the cell columns within the sub-placenta, analogous to cell columns of the human placenta [13]. Non-proliferative daughter cells leave the cell columns due to proliferation pressure, and start migrating in a self-secreted extracellular matrix [32]. Invasion of the maternal uterine tissue proceeds, stopped by either apoptosis or by the generation of multinucleated giant cells through polyploidization [33-35]. The patterns of trophoblast invasion in the guinea pig are similar to those shown in the degu placenta, where invaded maternal spiral arteries can be distinguished from non-invaded arteries by the presence of cytokeratin positive trophoblast versus SMA positive endothelium [29]. Previous investigations have focused on the use of immunohistochemistry to characterize the pattern of normal trophoblast invasion [12, 13, 29, 31, 36], but little is known about the molecular mechanisms, particularly in models of placental insufficiency.

In the present study, we show that MNR results in a reduction in the number of maternal vessels invaded by trophoblast and reduced trophoblast invasion at mid-pregnancy and is supported by the identification of decreased expression in transcripts involved in epithelial development and the regulation of cell migration. In particular, there was a 2.8-fold decrease in the expression of *Fgf10*. Fgf10 is a multifunctional mitogenic polypeptide that functions through ligand-receptor binding [37]. In humans, FGF10 is expressed in the extravillous trophoblasts in the first trimester and is known to activate ECM degradation pathways during trophoblast invasion [38]. Similarly, *Cxcl10*, was reduced in the MNR sub-placenta/decidua and has known roles in stimulating migration of various immune cells and modulating adhesion molecules [39]. Specific to placental development, CXCL10 is secreted by endometrial stromal cells [40] and expression is required for successful implantation through the regulation of trophoblast apposition and adhesion to the endometrium [41]. Hence, for the first time we have identified possible molecular mechanisms that are disrupted in placental insufficiency that result in a reduction in the invasive potential of trophoblast cells in the guinea pig MNR model, and that may be targeted in future therapeutic development.

Invasive trophoblast secrete metalloproteinases, metallopeptidases and endopeptidases in order to degrade the extracellular matrix [42, 43]. Protein expression of various metalloproteinases have been shown in the guinea pig sub-placenta/decidua [44]. Spatio-temporal changes in Mmp2 and Mmp9 expression across gestation suggest a functional role in relation to trophoblast invasion and placental angiogenesis [44]. Treatment of the placenta with the *hIGF1* nanoparticle did not change the pattern of trophoblast invasion in either the Control or MNR sub-placenta/decidua. However, this is unsurprising given the short time period between treatment administration and tissue collection. *hIGF1* nanoparticle treatment did however increase transporter and metallopeptidase activity in the MNR + *hIGF1* sub-placenta/decidua compared to MNR. Increased gene expression included *Spock1* which has previously been shown to be upregulated in tumors and promotes invasion [45], and various *AdamTS* homologues, which have previously been shown to modulate placental trophoblast invasion [46]. Nitric oxide synthase 1 (*Nos1*), which is responsible for synthesizing one of the most important vasodilators nitric oxide, expression was also increased in the MNR + *hIGF1* sub-placenta/decidua following *hIGF1* nanoparticle treatment. Together, these results indicate a potential to enhance trophoblast invasion in the MNR placenta. Further investigations extending the *hIGF1* nanoparticle treatment period and assessing trophoblast invasion are worthy of exploration, but beyond the scope of the current study.

Currently, the development of FGR cannot be predicted, and diagnosis is often made based on ultrasound measures which do not detect between 50-90% of cases [47, 48]. Therefore, a therapeutic intervention to treat FGR *in utero* must be safe to administer under situations where misdiagnosis occurs. In the current study pathway enrichment analysis indicated decreased expression of genes involved in kinase activity, and increased expression of genes relating to proteolysis. Such responses represent a downregulation in signaling mechanisms in order to maintain homeostasis in agreement with our previous investigations showing reduced ERK activity, and increased protein expression of the mTOR inhibitor DEPTOR, in the Control + *hIGF1* sub-placenta/decidua [30]. IGF1 elicits functional effects in placental trophoblast through binding with the IGF1 receptor; expression of which was reduced in the Control + *hIGF1* sub-placenta/decidua compared to Control. Ligand-receptor binding initiates signaling cascades including ERK/MAPK signaling and AKT/mTOR signaling [49], to modify gene expression. Our previous study [30] indicated reduced ERK phosphorylation was associated with reduced growth factor mRNA expression, including *VegfA* and *Pgf*, and is further supported by our current study which shows reduced expression of *Igf2, Kdr*, and *Nos2*.

In conclusion, the present study advances our understanding of the mechanisms contributing to placental insufficiency with a focus on trophoblast invasion in the middle of pregnancy. Using the developmentally relevant guinea pig model we identified changes to the transcriptome that likely result in the inadequate trophoblast invasion confirmed with morphological assessment. Additionally, this study has increased our understanding of pathways that *hIGF1* nanoparticle treatment acts on in order to restore or maintain appropriate placenta function. Overall supporting continual development of this technology in the correction of placental insufficiency and prevention of FGR.

## Statements and Declarations

### Competing Interests

Authors declare no conflicts of interest.

## Acknowledgments

We would like to thank Drs Craig Duvall and Mukesh Gupta for providing the co-polymer, Mrs. Kristin Lampe for assistance collecting placenta samples, and Dr Jason Puglise for assistance with RNA extractions. Additionally, we thank the University of Florida Interdisciplinary Center for Biotechnology Research Sequencing Core Facility for performing the RNA sequencing and Dr Alberto Riva for the bioinformatic analysis.

## Contributions

HNJ obtained funding, conceived the study and edited manuscript. BND performed experiments, analyzed data and edited manuscript. RLW conceived the study, performed experiments, analyzed data and wrote manuscript.

## Ethics approval

Animal care and usage was approved by the Institutional Animal Care and Use Committee at Cincinnati Children’s Hospital and Medical Center (Protocol number 2017-0065).

## Competing Interests

The authors have declared that no competing interest exists

## Data availability

All data needed to evaluate the conclusions in the paper are present in the paper and/or the Supplementary Materials. RNA Sequencing data has been uploaded to NCBI Geo under the accession number GSE269098.

## Funding

This study was funded by Eunice Kennedy Shriver National Institute of Child Health and Human Development (NICHD) award R01HD090657 (HNJ).

## Notes

### Competing Interest Statement

The authors have declared no competing interest.

